# Cellular and molecular mechanisms that shape the development and evolution of tail vertebral proportion in mice and jerboas

**DOI:** 10.1101/2024.10.25.620311

**Authors:** Ceri J. Weber, Alexander J. Weitzel, Alexander Y. Liu, Erica G. Gacasan, Robert L. Sah, Kimberly L. Cooper

**Affiliations:** Department of Cell and Developmental Biology, University of California, San Diego, 9500 Gilman Drive, La Jolla, CA 92093, USA; Shu Chien-Gene Lay Department of Bioengineering, University of California San Diego, La Jolla, California, USA

## Abstract

Despite the functional importance of the vertebral skeleton, little is known about how individual vertebrae elongate or achieve disproportionate lengths as in the giraffe neck. Rodent tails are an abundantly diverse and more tractable system to understand mechanisms of vertebral growth and proportion. In many rodents, disproportionately long mid-tail vertebrae form a ‘crescendo-decrescendo’ of lengths in the tail series. In bipedal jerboas, these vertebrae grow exceptionally long such that the adult tail is 1.5x the length of a mouse tail, relative to body length, with four fewer vertebrae. How do vertebrae with the same regional identity elongate differently from their neighbors to establish and diversify adult proportion? Here, we find that vertebral lengths are largely determined by differences in growth cartilage height and the number of cells progressing through endochondral ossification. Hypertrophic chondrocyte size, a major contributor to differential elongation in mammal limb bones, differs only in the longest jerboa mid-tail vertebrae where they are exceptionally large. To uncover candidate molecular mechanisms of disproportionate vertebral growth, we performed intersectional RNA-Seq of mouse and jerboa tail vertebrae with similar and disproportionate elongation rates. Many regulators of posterior axial identity and endochondral elongation are disproportionately differentially expressed in jerboa vertebrae. Among these, the inhibitory natriuretic peptide receptor C (NPR3) appears in multiple studies of rodent and human skeletal proportion suggesting it refines local growth rates broadly in the skeleton and broadly in mammals. Consistent with this hypothesis, NPR3 loss of function mice have abnormal tail and limb proportions. Therefore, in addition to genetic components of the complex process of vertebral evolution, these studies reveal fundamental mechanisms of skeletal growth and proportion.

## INTRODUCTION

Mammal skeletal diversity is remarkable. Much attention is paid to differences in the limbs and skull bones, but the vertebral skeleton is also strikingly different between species. For example, humans, dolphins, and giraffes all have seven cervical vertebrae but very different neck lengths^1,2^. In buffalo, extremely elongated neural spines extend from the dorsal side of thoracic vertebrae and serve as muscle attachment sites to support its massive head. At the far opposite end of the axial skeleton, tails range from externally absent in hominoid primates to elongate and prehensile adaptive appendages in new world monkeys^3^. How do the differences between vertebral size and shape develop and evolve?

Different parts of the vertebrate skeleton have distinct embryonic origins. The skull derives from neural crest and cephalic and somitic mesoderm^4^, the limbs emerge from lateral plate-derived buds^5^, and the vertebral/axial skeleton develops from the embryonic somites^6,7^. A substantial body of research has revealed the mechanisms of somitogenesis, which pinches off blocks of tissue in the trunk and tail region. A molecular clock determines the cadence of somite formation^8–11^, and maintenance of a progenitor pool determines the number of somites and thus the number of vertebrae, forming snakes in the most extreme cases^12,13^.

As the metameric series of somites march down the body axis, they acquire regional identities by translating their anterior-posterior position into expression of a ‘*Hox* code’^14–16^. *Hox* genes are anatomically expressed in reverse collinear order of their appearance in the genome with 3’ *Hox* genes in the anterior somites and 5’ Hox genes appearing posteriorly. These genes are both necessary and sufficient to define regional identities for groups of cervical (neck), thoracic (rib cage), lumbar (lower back), sacral (pelvic), and caudal (tail) vertebrae. For example, loss of function of the entire *Hox10* paralogous group results in lumbar vertebrae taking on a thoracic morphology with rib processes projecting from each element^17^.

However, once somites transform into cartilaginous vertebral scaffolds that will later become ossified bone, little is known about how they then acquire distinct shapes and sizes. Individual vertebrae are composed of a centrum with transverse, dorsal (neural), and/or ventral (hemal) processes^7^. The processes form articulations between neighboring vertebrae and between vertebrae and ribs, attachment sites for muscles, and protective structures for the spinal cord or dorsal artery and give distinctive shapes to each vertebra. Despite the relatively simple geometry of the centrum, the cylindrical core that lines up in series down the axis, its length also differs substantially between axial regions and even between neighboring vertebrae in series^18^.

The ossified vertebral skeleton not only has a distinct embryonic origin, but also predates the limb skeleton in its evolutionary origin by at least 60 million years^19–21^. Despite these differences, elongation of both limb bones and vertebrae occurs by a process of endochondral ossification of growth cartilages. However, while advances have been made to understand the cellular and molecular mechanisms that drive differences in limb bone elongation, far less is known of the mechanisms that control differential growth of the vertebral centra. How similar or different are the cellular and molecular mechanisms that control proportion in the limb and vertebral skeleton? How is skeletal modularity encoded in the genome such that individual bones can elongate and evolve independent of one another?

To answer these questions, we utilize the bipedal lesser Egyptian jerboa (*Jaculus jaculus*) as a model of extreme musculoskeletal proportion compared to quadrupedal laboratory mice. Thus far, much of our attention has focused on extreme elongation of the hindlimb and particularly the disproportionately long feet. However, the jerboa also has a disproportionately long tail, which is approximately 1.5x longer than the mouse normalized to body length^22^. Surprisingly, the long jerboa tail has three to four fewer vertebrae than in mice; their long tails are acquired by far greater elongation of individual vertebral elements in the mid-tail region. This substantially exaggerates the ‘crescendo and decrescendo’ of vertebral proportions that has been reported in the tail series of rodents, primates, and carnivores^3,18,23–26^.

Here we use the jerboa and mouse laboratory models to understand the temporal growth dynamics that establish adult vertebral proportion, the cellular drivers of differential growth, and candidate molecular mechanisms that determine and diversify vertebral proportion. We find that a greater number of chondrocytes undergoing endochondral ossification primarily drives differential elongation of the centrum, which is amplified by greater chondrocyte hypertrophy in the case of extremely long jerboa vertebrae. Intersectional differential expression analyses revealed 1,454 genes are disproportionately differentially expressed in the rapidly elongating jerboa 6th tail vertebra compared to shorter vertebrae in mouse and jerboa. Most of these (80.3%) are not disproportionately differentially expressed in studies of limb bone proportion, but there is significant overlap with genes found in each limb proportion study suggesting a subset of mechanisms do control proportion throughout the skeleton. Among these, natriuretic peptide receptor C (NPR3) appears in similar studies of jerboa, mouse, and rat limb proportion and a GWAS analysis of human body proportion^27–29^. Perturbation of this signaling pathway in the mouse disrupts tail vertebral proportion, suggesting that this pathway may be a crucial regulator of proportion in the vertebral skeleton in addition to its previously alluded to regulation of limb proportion. Our work has therefore extended observations of differences between the tail skeletons of adult mice and jerboas to identification of cellular and molecular mechanisms that establish vertebral proportion and lays a foundation to uncover additional mechanisms of skeletal modularity that shape the extraordinary diversity of mammals.

## RESULTS

### Development of adult tail proportion

We previously reported that the adult jerboa tail is 1.5x the length of a mouse tail, normalized to body length, with three to four fewer vertebrae^22^; vertebral counts vary by one in both species (Fig. 1A-B). To achieve this, the vertebrae in the jerboa mid-tail region reach far greater lengths to disproportionately increase overall tail length relative to body length. To identify when differences in overall tail length and vertebral proportion within the tail first appear and how they manifest, we collected mice and jerboas at weekly intervals from birth to mature tail proportion at six weeks of age (postnatal day 42; P42). We measured the ratio of overall tail length to body length (naso-anal distance) and the lengths of each ossified tail vertebral centrum (diaphysis) in the anterior-posterior axis from micro-computed tomography (µCT) images (Fig. 1).

**Figure 1.**
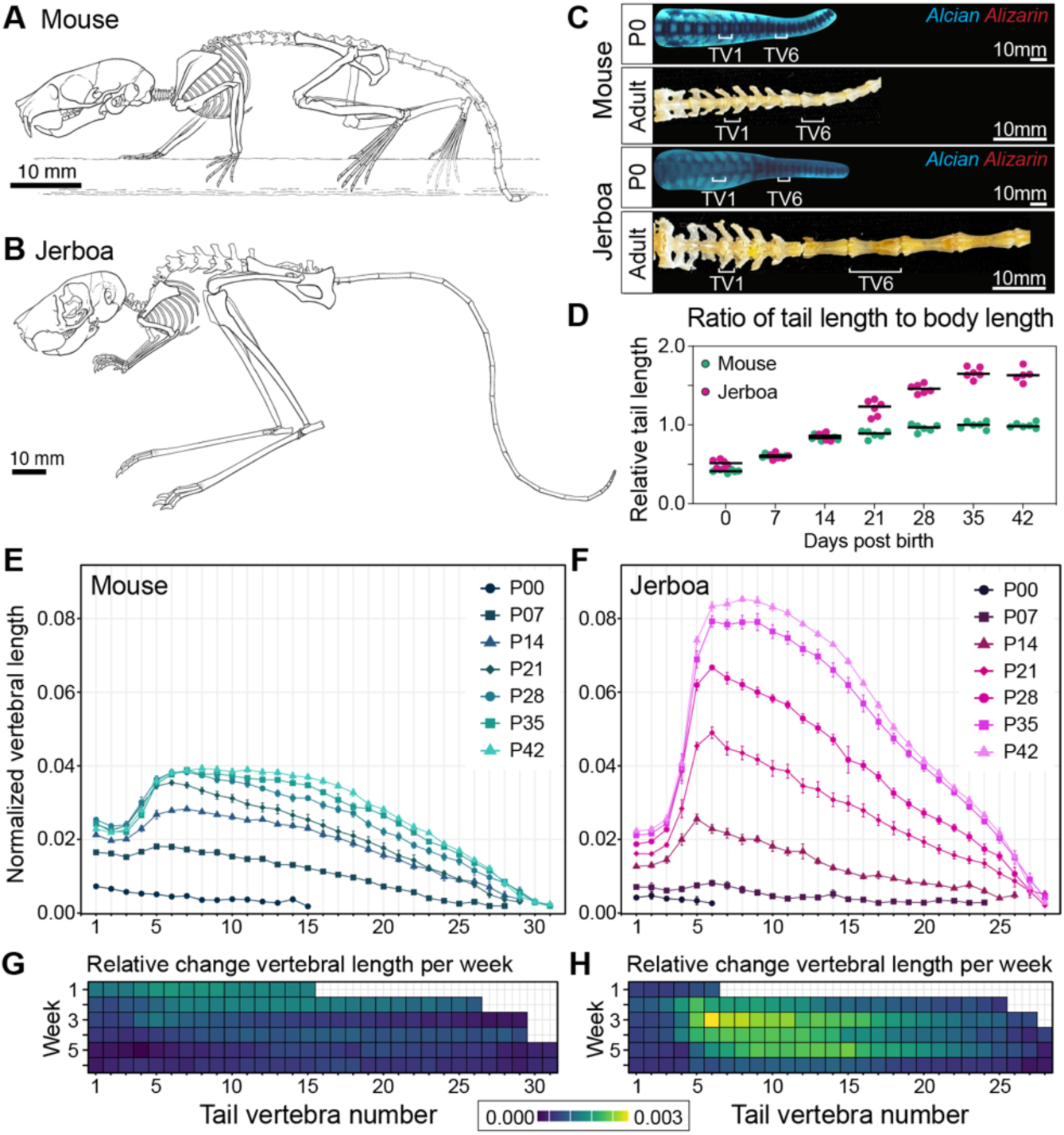
Development of mouse and jerboa tail proportion. (A-B) Diagram of adult mouse (A) and jerboa (B) skeletons modified from Moore et al. 2015. (C) Alcian and alizarin-stained neonates show all proximal vertebral elements are present at P0 in both species. Cleaned adult proximal tail skeletons show similar vertebral morphologies despite differences in size and proportion. (D) Mouse and jerboa tails are about half of the naso-anal length at birth. Tail proportion diverges by P21; the mouse tail remains about equal to body length while the jerboa tail elongates to 1.5x the body length. (E-H) Lengths of vertebral centra measured weekly from birth to six weeks, normalized to the naso-anal length of each mouse (E) and jerboa (F). The weekly relative change in length of each vertebra is represented in a heat map. The greatest rate of change is yellow and least in dark blue with the scale equivalent for mouse (G) and jerboa (H).

At birth, the normalized lengths of mouse and jerboa tails are about half of the naso-anal distance (Fig1. C-D, Fig. S1), and individual vertebrae are sequentially shorter from the first tail vertebra to the tip of the tail in both species (Fig. 1E-F). Interestingly, although both species have formed all the cartilaginous vertebral scaffolds by birth, 31 in mice and 28 in jerboas (n=2 each), only half as many tail vertebrae are ossified in jerboas compared to mice (Fig. 1C, 1E-F). A similar delay in ossification has been observed associated with extreme elongation of jerboa metatarsals and bat metacarpals^30–32^.

These data suggest that the difference in total tail length and vertebral proportion, within and between species, does not emerge during somitogenesis or early chondrogenesis but rather during postnatal endochondral ossification of the centra. From birth to postnatal day 14 (P14), the overall tail length to body length ratio maintains similar proportion in the two species; tail and body elongation are evolutionarily isometric for the first two weeks (Fig. 1E-F, Fig. S1). At postnatal day 7 in both mice and jerboas, a disproportionately rapid rate of elongation is initiated in the mid-tail region with a plateau of the greatest lengths centered on the 5^th^-8^th^ tail vertebrae (TV5-TV8) (Fig. 1E-H) with TV6 demonstrating the greatest change in vertebral length in each species. Vertebral lengths then sequentially decrease to the tail tip. This “crescendo-decrescendo” is amplified by continued postnatal differential growth, far more rapidly in jerboas, and adult tail proportion is achieved in both species by the fifth week (Fig. 1E-F). In jerboas, the most rapid disproportionate elongation of mid-tail vertebrae occurs between P14 and P21, the time at which the overall tail length to body ratio also begins to diverge from mouse (Fig. 1D, 1H).

Interestingly, this period of the most rapid vertebral elongation occurs one week later in jerboas than in mice, consistent with our observation that the initial ossification in the tail series, and formation of the secondary ossification centers (vertebral endplates) is also delayed in jerboas compared to mice (Fig. S2). This supports the hypothesis that disproportionately rapidly elongating skeletal elements in jerboa behave as “younger” growth cartilages enabling faster growth for longer than homologous elements of mice^30^.

### Cellular parameters of tail vertebral elongation

Each vertebral centrum elongates by endochondral ossification of the cranial and caudal growth cartilages, though not necessarily at the same symmetric rate. To quantify the relative rates of growth contributing to vertebral elongation, we focused on the vertebrae with the most similar (TV1) and most different rates of elongation (TV6) between species in the proximal to mid-tail (Fig. 2A-C). We selected a timepoint within the window of greatest disproportionate elongation for each species prior to formation of the secondary ossification centers, P5-7 for mouse and P14- 16 for jerboa. To quantify the daily rate of growth cartilage elongation during this growth phase, we injected pups with a pulse of calcein dye to fluorescently label mineralized bone (at P5 for mouse and P14 for jerboa), two days prior to collection at P7 or P16 (Fig. S3). The distance between the calcein-labeled bone and the chondro-osseus junction of the growth cartilage, divided by two, reveals the growth rate as the length of new bone formed per 24-hour period.

**Figure 2.**
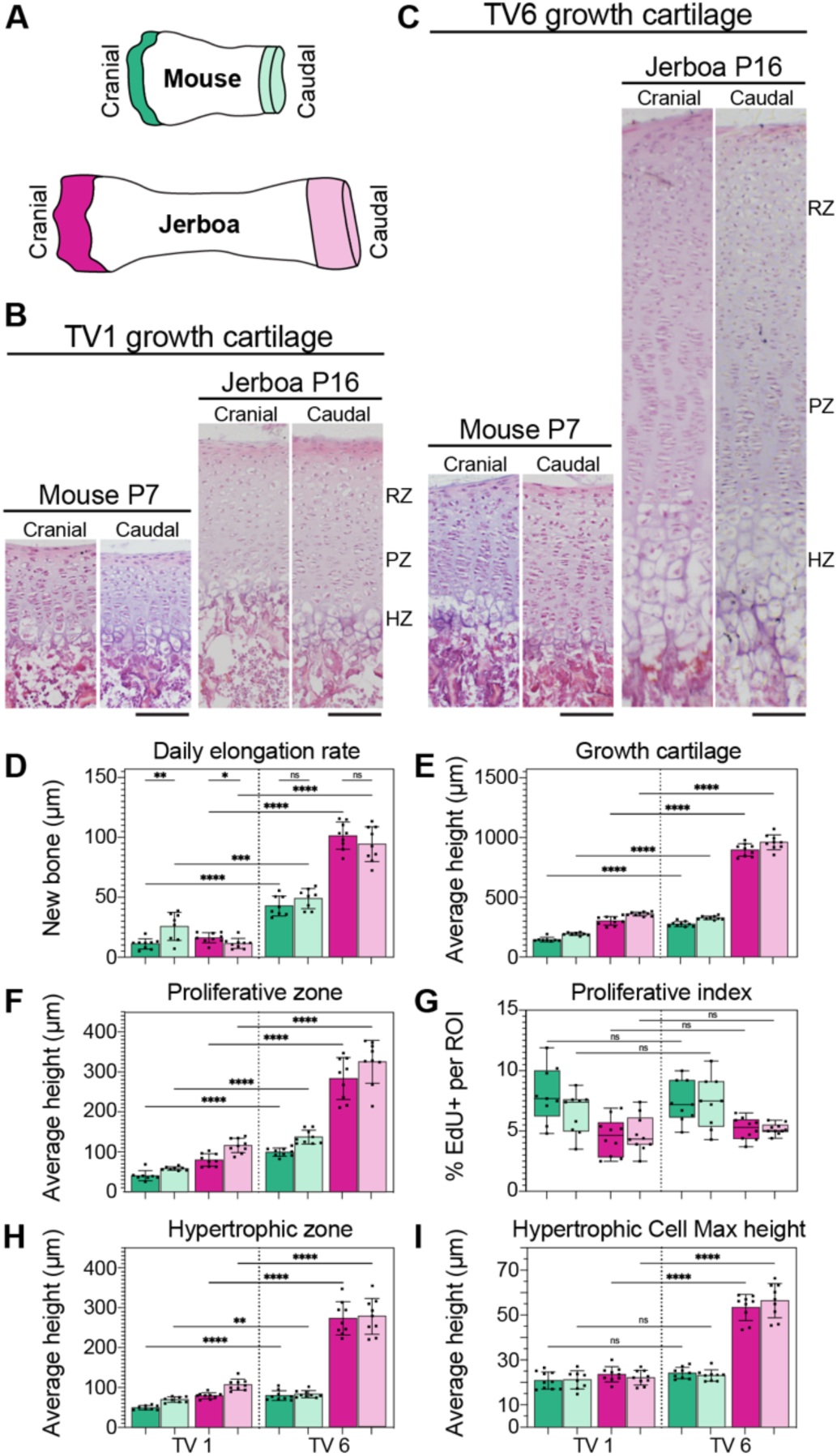
Cellular parameters of growth during the greatest difference in vertebral elongation rate. (A) Schematic of tail vertebrae with cranial and caudal growth cartilage color schemes used throughout. (B-C) Histology of mouse and jerboa cranial and caudal growth cartilages of TV1 (B) and TV6 (C) during rapid elongation. (RZ=resting zone, PZ=proliferative zone, HZ=hypertrophic zone). (D-I) Growth cartilage parameters in each cranial and caudal of mouse and jerboa TV1 and TV6. (D) Daily elongation rate. (E) Height of the growth plate. (F) Height of the proliferative zone. (G) Proliferative index calculated as the fraction of EdU+ cells of all cells in an ROI. (H) Height of the hypertrophic zone. (I) Maximum height of hypertrophic chondrocytes in the direction of bone elongation. Welch’s t-test, n=>9 of mixed male and female animals, * ≤ 0.05, ** ≤ 0.01, *** ≤ 0.001, **** ≤ 0.0001.

During this window of greatest differential growth of TV6 versus TV1, we found that cranial growth cartilage elongation is significantly slower than caudal elongation within mouse TV1. Conversely, the cranial growth cartilage elongates slightly but significantly faster compared to the caudal growth cartilage in jerboa TV1 (Fig. 2D, Fig. S3). There is no significant difference between the cranial and caudal growth rates of TV6 in either species. However, consistent with µCT measurements of whole vertebrae, both cranial and caudal growth cartilages elongate significantly faster in TV6 than in TV1 in both species, and the jerboa TV6 growth cartilages elongate more than 2x faster than mouse.

In the limb skeleton, multiple chondrocyte parameters (e.g., the number of chondrocytes produced each day, the volume of extracellular matrix, and chondrocyte hypertrophy) all differ between growth cartilages that elongate at different rates^33–35^. In reptiles and birds, differential growth primarily correlates with the number of cells in the growth cartilage, particularly the height of the proliferative zone^36,37^. However, growth rate differences in mammal limbs, within and between species, are additionally driven by the final hypertrophic cell size^31–33,35,38–40^. The doubling of an individual flattened proliferative chondrocyte adds just 8-9 µm to the axis of elongation whereas each of these cells adds up to 40-50 µm to the growth axis as it proceeds through hypertrophic enlargement^31,38^. Hypertrophic cell sizes also differ markedly between mammal limb bone growth cartilages and contribute the most to differences in the daily rate of elongation^38^. We therefore measured growth cartilage height, proliferative zone height and proliferation index, and hypertrophic zone height and maximum hypertrophic chondrocyte size to distinguish relative cellular contributions to differential vertebral elongation in cranial and caudal growth cartilages of TV1 and TV6 in mouse and jerboa.

We first measured total growth cartilage height from the intervertebral joint surface to the chondro-osseous junction. In both species, growth cartilage heights are significantly greater in the more rapidly elongating TV6 than in TV1 (Fig. 2B-C, 2E; Fig. S3). Consistent with the greater than two-fold difference in growth rate, both cranial and caudal growth cartilages of jerboa TV6 are more than twice the height of any other measured growth cartilage. The heights of individual zones within the growth cartilage also correlate with growth rate (Fig. 2F, 2H, Fig. S3). A greater proliferative zone height could be due to faster cell-cycle time that produces more cells before they initiate hypertrophic differentiation, and/or a larger number of progenitor cells that double at the same rate. To test this, we measured the proliferation index within each growth cartilage during the rapid window of growth by injecting EdU into pups two hours before collection at P7 (mouse) or P16 (jerboa) to label cells in S-phase of the cell cycle. A greater fraction of cells in S-phase would indicate a more rapid rate of division. However, we found that the fraction of S-phase chondrocytes is not significantly different between TV6 and TV1 in either species (Fig. 2G, Fig. S4). This suggests that the difference in proliferative zone height is due to a greater number of chondrocytes dividing at a similar rather than a faster rate of progression through the cell cycle.

Consistent with this hypothesis that a greater number of chondrocytes progress through endochondral ossification of the most rapidly elongating growth cartilages, we found that the height of the hypertrophic zone also correlates with growth rate (Fig. 2H). Since hypertrophic chondrocyte size drives growth rate differences in mammal limb bones^31–33,35,38–40^, we also measured the average maximum diameter of these cells in the axis of elongation in cranial and caudal growth cartilages of mouse and jerboa TV1 and TV6. Hypertrophic cell size is only marginally or not significantly different in mouse growth cartilages and in jerboa TV1. However, the extreme rate of jerboa TV6 elongation is driven in part by hypertrophic chondrocytes that are more than twice the height of chondrocytes in the other vertebral growth cartilages (Fig. 2I). This was surprising, because even hypertrophic chondrocytes in the rapidly elongating jerboa metatarsus are only ∼58% larger than those in the mouse. Comparing limb and vertebral growth cartilages revealed that hypertrophic cell sizes are similar between jerboa TV6 and jerboa metatarsals^31^, and that the larger percent difference in the vertebrae is because other vertebral hypertrophic chondrocytes are much smaller than in limb bones. This suggests that hypertrophic chondrocyte size is evolvable in mammalian vertebral growth cartilages, but it might only be a driver of extreme vertebral proportion compared to differences between limb bones in a breadth of mammals.

Collectively, these data suggest that the tail crescendo-decrescendo is largely driven by differences in the number of cells progressing through endochondral ossification in each vertebra, as in bird and reptile limbs bones. This difference between vertebrae is greatly amplified in the jerboa due to exaggerated enlargement of the jerboa TV6 hypertrophic chondrocytes, driving extreme disproportionate elongation of the mid-tail. What are the molecular differences that may cause these cellular differences that drive the development and evolution of vertebral proportion?

### Interspecies intersectional transcriptomics reveal candidate molecular mechanisms of tail diversification

Multiple studies have investigated the genetic basis of skeletal proportion by identifying gene expression differences between limb bone growth cartilages^28,29,41^. Previously, we applied such an approach to identify candidate drivers of limb skeletal proportion in the jerboa, with its elongate hindlimb and disproportionately long feet compared to mice^29^. Compared to limb bone elongation, far less is known about the molecular mechanisms of vertebral elongation to establish skeletal proportion or to what extent these mechanisms are shared with long bones of the limb. Similar careful intersection of comparative gene expression studies can identify candidates for the modular regulation of vertebral elongation to establish axial proportion.

Because the cranial and caudal contributions to vertebral elongation are very similar in all growth cartilages, but the difference in relative contribution to growth is greatest between cranial growth cartilages of mouse TV1 and TV6 (Fig. 2D), we chose to focus on cranial growth cartilages for these analyses. We performed bulk-RNA sequencing on three pooled cartilages per each of four replicates during the windows of greatest disproportionate growth in mouse and jerboa, at P7 and P16 respectively, before the appearance of secondary centers of ossification (Fig. 3A, Fig. S2). To directly quantify differences in gene expression between jerboa and mouse homologous skeletal elements, we first annotated a set of 17,640 orthologous genes using TOGA and the *Mus musculus* (mm10) and revised *Jac jaculus* (mJacJac1.mat.Y.cur) genome assemblies (NCBI). We included both one-to-one orthologous transcripts and genes in the one-to-zero classification, because predicted functional loss might be explained by sequence and/or assembly errors.

For each of the four sample sets, mouse and jerboa TV1 and TV6, we sequenced four biological replicates and mapped reads to the respective genome using the TOGA orthologous gene set annotations. We then used DESeq2 to perform differential expression analyses within and between species with an additional normalization step to account for gene length differences between species^29,42^. Principal components analysis (PCA) revealed that the variation between samples is primarily due to species differences (PC1) and next by growth cartilage origin (PC2) (Fig. S5).

To narrow our data set to candidate genetic drivers of the evolution of tail proportion by differential rates of vertebral elongation, we performed a series of comparisons and intersections of differential expression datasets (Fig. 4A-C). Jerboa TV6 elongates at a disproportionately faster rate compared to mouse TV6, and TV1 is more similar in both species. We therefore directly compared expression in homologous vertebrae (jerboa TV6 to mouse TV6; jerboa TV1 to mouse TV1) (Fig. 4A). We reasoned that genes that drive the disproportionate elongation of jerboa TV6 should be disproportionately differentially expressed in jerboa TV6. These fall within two categories: 1,864 genes are significantly differentially expressed between jerboa and mouse TV6 (padj<0.05) but not between species in TV1 (Fig. 4E, Table S1). We then intersected the log_2_-fold change values of all genes that are significantly differentially expressed between species in both vertebrae (Table S2). A majority of these are near-equivalently differentially expressed (slope=0.883) and are therefore unlikely to contribute to differential elongation of jerboa TV6. Exclusion of genes within the 95% prediction interval of the correlation leaves 421 disproportionately differentially expressed genes in this subset, which we added to the 1,864 genes that significantly differ between species only in TV6 (Fig. 4D, Table S3).

**Figure 4.**
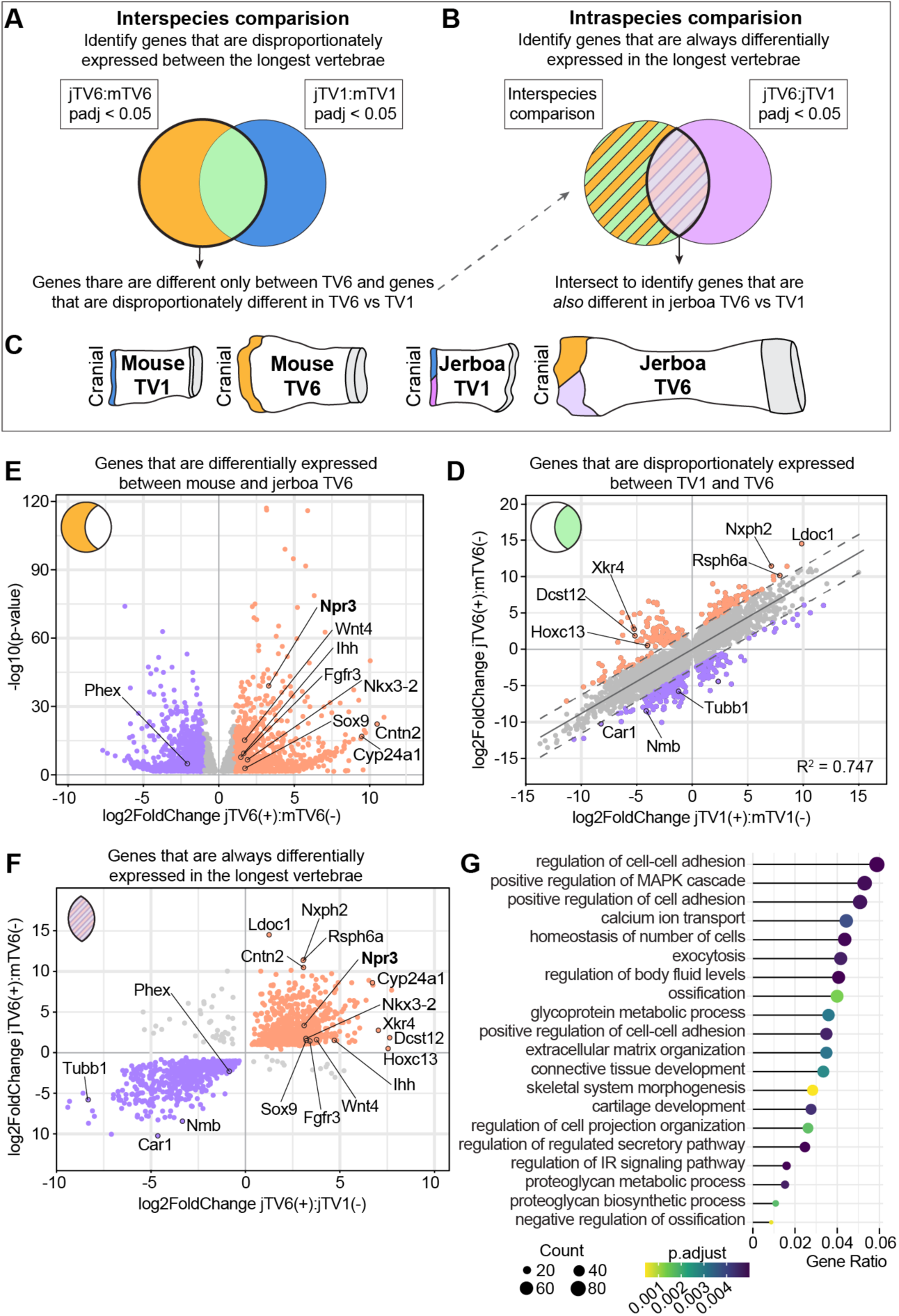
Design and analysis of intersectional interspecies transcriptomics. (A-C) Schematics of the interspecies and intraspecies comparison approaches of differential expression analyses. (E) 1,864 genes are significantly differentially expressed between mouse and jerboa TV6 but not between species in TV1. Genes that are expressed higher in jerboa TV6 are orange; lower are purple. (F) 6,786 genes are significantly differentially expressed between jerboa and mouse both in TV6 (y-axis) and in TV1 (x-axis); most are equivalently differentially expressed (gray points, slope=0.883), but 421 genes are outside of the 95% prediction interval and designated disproportionately differentially expressed. (F) Of all 2,285 genes highlighted in D and E, 1,454 are also differentially expressed in jerboa TV6 versus TV1, consistently in the same direction in the longest versus shortest element. Genes in orange are expressed higher in jerboa TV6 compared to mouse TV6 and in jerboa TV6 compared to jerboa TV1, while those lower in both comparisons are in purple. (G) Selection of GO terms relevant to cartilage that are enriched among the 1,454 candidate genes.

We then reasoned that genes that control the disproportionate rate of jerboa TV6 elongation should also differ in their expression between jerboa TV6 and jerboa TV1. Intra-species comparison within jerboa revealed 7,911 genes that are significantly differentially expressed between vertebrae despite sharing embryonic origins, tail regional identity in the axial skeleton, and developmental age (Fig. 4B, Table S4). We intersected these with disproportionately differentially expressed genes between species and found 1,454 that are significantly differentially expressed in both datasets with the sign value consistently correlating with the most rapid rate of elongation in jerboa TV6 (Fig. 4F, Table S5).

GO term enrichment analyses provide valuable insight into possible biological processes regulating disproportionate vertebral elongation, even though knowledge bases are restricted to the fraction of genes that have been studied in limited biological contexts^43^. To identify biological processes enriched in this set of 1,454 genes that are differentially expressed within jerboa and also disproportionately between species, we implemented the clusterProfiler package^44,45^. Among the top significantly enriched terms, we find many biological processes critical to cartilage development and ossification (Fig. 4G).

A major goal of this study was to determine if genes that control vertebral proportion overlap significantly with the genes that have been associated with limb bone proportion, which would suggest modular control of a global genetic ‘tool-kit’ of bone growth. In addition to our previous work that identified candidate mechanisms of disproportionate jerboa hindlimb elongation^29^, an expression study in mice and rats identified genes that are differentially expressed in growth cartilages that elongate at different rates in the same young animal (fast proximal tibia versus slower distal phalanx)^28^. The same study also identified genes that are differentially expressed in the rapidly elongating young tibia versus the older tibia that slows its growth rate. Although not an equivalent differential expression analysis, a genome wide association study identified single nucleotide polymorphisms in a human population that are associated with variance in the proportion of leg length with respect to crown-rump length^27^.

We intersected these four datasets (listed in Table S6) with our set of genes that are associated with the evolution of tail vertebral proportion. A Fisher’s exact test with Benjamini– Hochberg multiple hypothesis correction found that there is significant overlap between our gene set and all other datasets. Molecular mechanisms that establish vertebral proportion are therefore significantly similar to mechanisms that control limb bone proportion.

To identify genes with the strongest indication of a mechanistic role in the broad regulation of skeletal proportion, we collated genes that also have reported “short tail” or “long tail” mutant phenotypes in the Mouse Genome Informatics (MGI) database (Table S6)^73,74^. Of the 1,454 disproportionately differentially expressed genes that correlate with the rapid elongation of jerboa TV6, (Table S7), 20 of these have a reported tail mutant phenotype and/or appear in three or more limb proportion datasets. Many of these are known regulators of chondrocyte proliferation and/or maturation into hypertrophy. Although we cannot say at this time how many mutations affect the evolution of skeletal proportion, all of these genes are strong candidates, either by their own modular *cis*-regulatory control or by modular expression of upstream transcription factors.

A single gene, *Npr3*, is common to all five analyses (Table 1, S7). *Npr3* is more highly expressed in the rapidly elongating jerboa TV6 compared to mouse TV6 and jerboa TV1, higher in jerboa metatarsals than in mouse^75^, and higher in the most rapidly elongating mouse and rat limb bones by age or by location^28^. These data suggest that *Npr3* may be a major contributor to the development and evolution of proportion broadly in the skeleton.

**Table 1.**
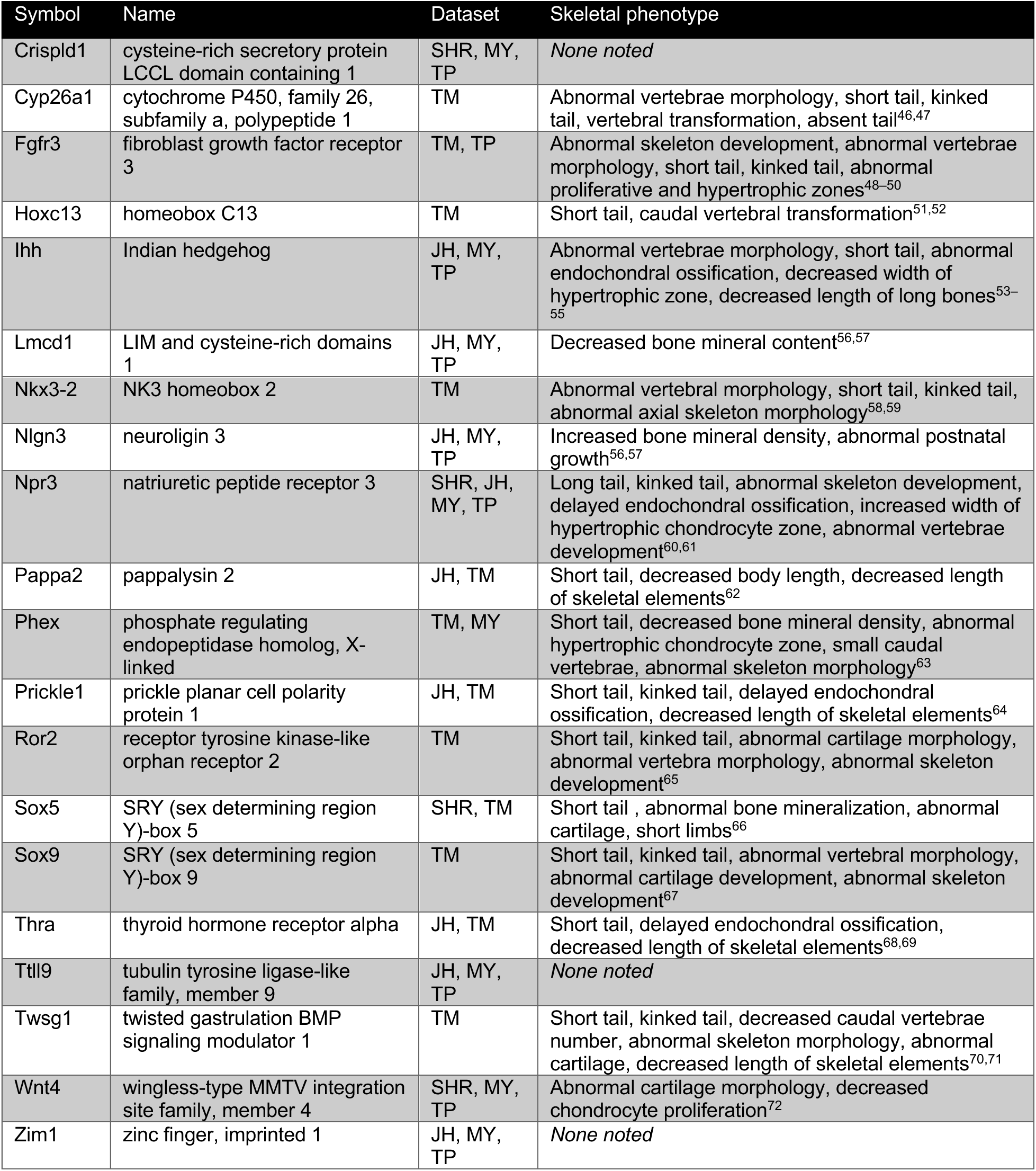
20 candidate genes associated with vertebral and limb proportion. Mutant phenotypes identified in MGI were confirmed in primary literature. <u>JH</u> – candidate genes driving <u>J</u>erboa <u>H</u>indlimb elongation^26^ <u>MY</u> – genes that are differentially expressed in the rapidly elongating <u>Y</u>oung tibia versus the <u>M</u>ature tibia^25^ <u>TP</u> – genes that are differentially expressed in the fast-growing proximal <u>T</u>ibia versus slow distal <u>P</u>halanx ^25^ <u>SHR</u> – SNPs associated with variance in the proportion of leg length with respect to total body length^30^ <u>TM</u> – genes with reported “short <u>T</u>ail” or “long <u>T</u>ail” <u>M</u>utant phenotypes in the MGI database^31,32^.

### Regulated natriuretic peptide signaling contributes to tail vertebral proportion

NPR3 belongs to a family of receptors that bind c-type natriuretic peptide (CNP) and regulate diverse cellular processes from cardiac function and blood pressure to endochondral ossification^76–78^. NPR2, the primary signaling receptor of this family, is expressed in pre-hypertrophic chondrocytes and is crucial to normal bone elongation by regulating MAPK signaling through cGMP-mediated activation of PKG^79–81^. Natriuretic peptide receptor 3 (NPR3) is an inhibitory regulator of natriuretic peptide signaling through clearance mediated degradation of the natriuretic peptide ligand^60^. Mutations in the NPR3 receptor typically cause skeletal overgrowth, presumably through enhanced activation of the NPR2 receptor due to excess available natriuretic peptide^60,61,80,82^. Loss of NPR2 and CNP (*Nppc*) expression in humans causes Acromesomelic Dysplasia Maroteaux Type (AMDM), distinguished by disproportionate shortening of the most distal limb elements, implicating NPR signaling in establishing proportion in the limb skeleton^83–90^. However, while inactivating mutations in *Npr2* and *Npr3* have been reported to shorten or lengthen the mouse tail respectively, consistent with overall effects on body size, alterations to tail or vertebral proportion were not documented^60,61,77,83,91,92^. We therefore obtained *Npr3* knock-out mice to investigate the effect of loss of NPR3 on overall tail-to-body and individual vertebral proportions.

We collected *Npr3^-/-^* mice and wild-type littermates once they reached adult proportions at postnatal day 42 (6 weeks) and prepared the sacral and caudal tail skeleton for µCT analysis. Compared to mice from our temporal analysis of typical tail growth (CD1 genetic background), total tail length to naso-anal length is not significantly different in wild-type siblings (C57/Bl6 genetic background) (Fig. 5A). However, *Npr3^-/-^* mice have a significantly longer tail to body length ratio, though they also have characteristic but variable spinal kyphosis by this age (Fig. 5A). We then measured lengths of individual tail vertebral centra in *Npr3^-/-^*and wildtype siblings and compared these normalized lengths to our CD1 P42 vertebral measurements (Fig. 5B). Although the scan resolution of *Npr3^-/-^*mice and littermate controls differs compared with our time course data (50 µm^3^ versus 9 µm^3^), which could contribute to absolute differences, the ‘crescendo-decrescendo’ curve and individual vertebral lengths are similar between wild-type siblings and CD1 mice. By contrast, the proximal and mid-tail vertebrae are disproportionately longer in *Npr3^-/-^* mice, whereas individual vertebral lengths are not significantly different from TV13 to the tail tip. We speculate that the longer proximal vertebral elements may explain the ‘typical cone-shaped implantation of the tail’ that was previously documented in *Npr3^-/-^* mice^61^.

**Figure 5.**
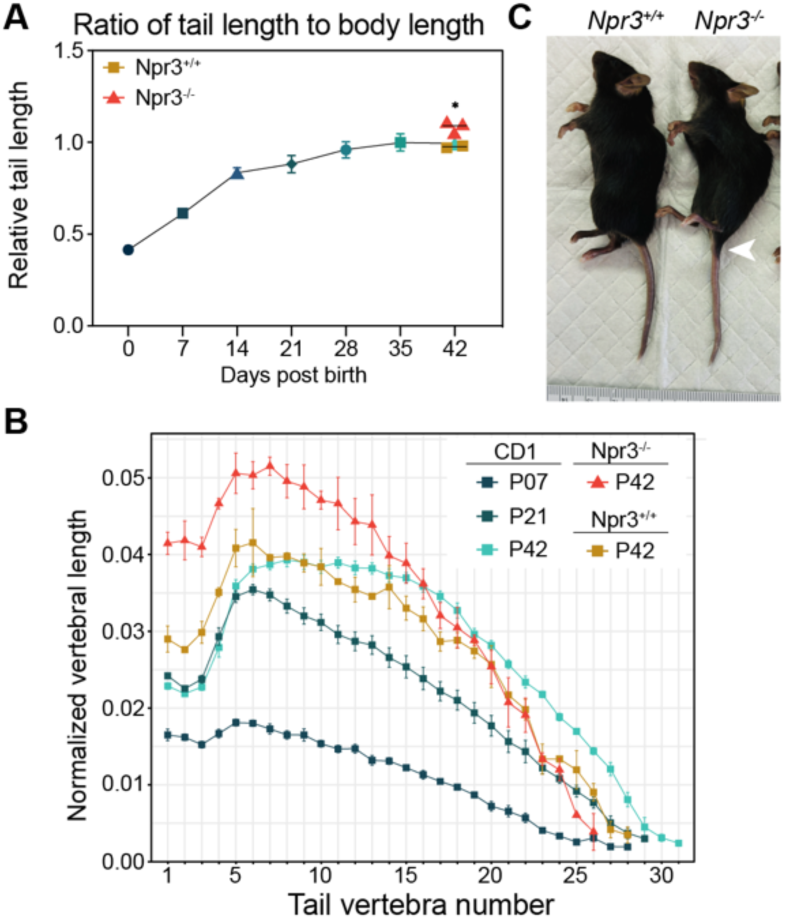
Natriuretic peptide signaling affects tail vertebrae proportion. (A) Ratio of tail length to naso-anal distance in adult (P42) *Npr3^-/-^*mice (orange triangle) and wildtype *Npr3^+/+^* siblings (gold square) compared to average relative tail lengths in CD1 mice (shades of blue). (B) Lengths of vertebral centra at P42 normalized to the naso-anal length of each *Npr3^-/-^* (orange triangle) and wildtype *Npr3^+/+^* siblings (gold square). Normalized vertebral lengths for CD1 mice at P07, P21, and P42 are displayed for comparison. The distal-most vertebrae in all *Npr3^-/-^*mice and siblings were not measurable due to the lower µCT-scanner resolution. (C) *Npr3^-/-^* mouse with wildtype sibling aged P42. White arrowhead points to ‘cone-shaped implantation of the tail’.

## DISCUSSION

Here we used the remarkable diversity of the axial skeleton, both within an individual and between species, to study how serially repeating elements achieve distinctly different sizes. We provide valuable insights into how the axial skeleton establishes different elongation rates between vertebrae considered to have the same regional “identity”. Our intersectional approach, comparing vertebrae with disproportionate (TV6) to those with similar growth rates (TV1), allowed us to identify cellular mechanisms and associated candidate genes for control of disproportionate vertebral elongation.

Mammal tails differ in both the number and lengths of individual vertebrae^3,25,26,93^. In addition to differences in the developmental mechanisms that determine the number of vertebrae and their relative lengths, the two characters are also genetically separable. In deer mice (*Peromyscus maniculatus*), six genomic regions (loci) contribute to tail length differences between forest and prairie ecotypes. Three of these loci contribute to differences in vertebral number but not length and three to length but not number^26^.

Research in laboratory models (mice and chickens) has shown that somitogenesis to determine vertebral number is controlled in part by genes in a *Gdf11/Lin28/Hox13* interaction network^94,95^. Knowledge of this network focused attention on *HoxD13* among 168 genes in the three deer mouse ‘vertebral number’ loci, and further investigation found that differences in the non-coding control of *HoxD13* expression likely contribute directly to the difference in number of deer mouse tail vertebrae^25,26^. However, details of the potentially hundreds of genes in the ‘vertebral length’ loci remain unreported, largely because less was known to winnow the list down to likely functional candidates. Our analyses of the cellular and molecular differences driving differential growth within and between jerboas and mice is therefore a crucial first step to identify causative mechanisms of vertebral length diversification.

Despite the sizable gap in knowledge of the mechanisms of vertebral elongation, we do have a rich understanding of bone growth from decades of studies largely focused on the limb skeleton. However, even though both limb and vertebral elements elongate by endochondral ossification of a growth cartilage, the appendicular and axial skeletons have distinct developmental origins. Furthermore, the axial endochondral skeleton likely predates tetrapod limbs by at least 60 million years^19–21^. It was therefore entirely unclear whether the cellular and molecular mechanisms of differential growth to establish and vary proportion should be similar or distinct.

Growth cartilage height, or the number of chondrocytes that proceed through proliferation and differentiation, corresponds strongly with growth rate in bird, reptile, and mammal limb bones; we show here that growth cartilage height also associates with tail vertebral growth rate in mice and jerboas. In mammal limb bones, however, hypertrophic chondrocyte size also differs significantly in growth cartilages that elongate at different rates and is indeed the greatest contribution to differences in the daily rate of limb bone elongation within and between many mammal species^31–33,35,38–40^. By contrast, we show that hypertrophic chondrocyte cell size does not differ in rapidly elongating mouse sixth vertebral cartilages compared to the slower-elongating first vertebrae, and these are smaller than limb bone hypertrophic chondrocytes. Surprisingly, however, the extremely rapidly elongating sixth vertebra in the jerboa tail has hypertrophic chondrocytes that are around twice the size of other vertebral hypertrophic chondrocytes, about four times the difference in cell size between rapidly elongating jerboa metatarsals compared to mice^31^. Together, this suggests that growth cartilage height and cell number is readily evolvable in the limb and vertebral skeleton, possibly traceable to the origin of tetrapods, whereas hypertrophic cell size differs in mammal limbs and in an extreme case of vertebral elongation.

There are also striking similarities and differences between limb and vertebral proportion at a molecular level. Here, we show significant overlap of differentially expressed genes in our vertebral comparisons, three limb differential expression datasets, and a human GWAS analysis of limb to trunk proportion suggesting some of the same mechanisms control proportion in the limbs and vertebrae. However, despite the significance of overlap, a majority (80.9%) of the genes that are differentially expressed between tail vertebrae are not among limb proportion genes, and genes among these might therefore control vertebral proportion independent of the limb skeleton. One of these genes, *HoxC13,* is a noted regulator of posterior embryonic development and segment identity^52^. Intriguingly, loss of *HoxC13* also extends the shapes of vertebrae, typical to TV4-6 (transverse processes, see Fig.1C), distal to the levels of TV9-11, and the longest vertebra shifts together with this ‘homeotic transformation; conversely, *HoxC13* overexpression reduces tail length^51,52^.

Like many of the genes that are differentially expressed only between vertebral elements and not within limbs, *Nkx3-2* does play an important role in bone growth throughout the skeleton*. Nkx3-2* partners with *Sox9* to initiate chondrogenesis in the somites; both are later primarily expressed in resting and proliferating chondrocyte zones^96–99^. In limbs, *Nkx3-2* has been shown to promote chondrocyte survival and proliferation and also delays chondrocyte maturation through repression of *Runx2*^96^. *Nkx3-2* was also identified in a genomic analysis after artificial selection for standing variation in mice drove disproportionate elongation of the tibia, though the proposed *cis*-regulatory mutations may lower expression of *Nkx3-2*^41^. Similar to our study, tibia elongation was also achieved by increasing the number of chondrocytes going through endochondral ossification, but *Nkx3-2* enhancer activity was reduced by selection for a longer tibia^41^. In jerboa TV6 when growth is the fastest, higher *Nkx3-2* might expand the growth cartilage by both stimulating proliferation and delaying maturation of those chondrocytes.

Also shared among vertebral and limb datasets, we find regulators of chondrocyte maturation that are crucial throughout the skeleton, including *Ihh, Wnt4,* and *Fgfr3. Ihh* regulates chondrocyte proliferation and hypertrophy in a negative feedback loop with PTHrP, promoting proliferation and maintaining bone growth while indirectly delaying hypertrophy^97,100–103^. In contrast, *Wnt4* stimulates chondrocyte hypertrophy, and overexpression of this gene reduces chondrocyte proliferation, because maturation is accelerated^72,104^. During postnatal growth, *Fgfr3* is expressed in the proliferative zone and suppresses chondrocyte proliferation and hypertrophy via activation of STAT and MAPK/ERK signaling respectively^105–108^. Identification of these genes in our final candidate list suggests that the central components of the chondrocyte maturation process are locally tuned to regulate the development and evolution of limb and vertebral proportion.

Activating mutations in the *Fgfr3* receptor are the most common cause of human achondroplasia^109^. Because of pathway interactions between FGF and natriuretic peptide receptor (NPR) signaling^78,80,81^, Vosoritide, a c-type natriuretic peptide analog, is a relatively new intervention to treat complex aspects of the syndrome^81,110,111^. Interestingly, natriuretic peptide receptor C (*Npr3*) is the only gene we found in all datasets of limb and vertebral differential expression, human body proportion GWAS, and mouse tail mutant phenotypes suggesting it plays a crucial role in establishing and diversifying skeletal proportion.

Paradoxically, although NPR3 loss-of-function mice have disproportionately longer tails with disproportionately longer mid-tail vertebrae, NPR3 expression is consistently *higher* in the fastest growing limb bones of mice, rats, and jerboas as well as jerboa TV6 compared to bones that elongate at a slower rate. NPR3 is thought to suppress NPR2-mediated promotion of bone elongation. NPR2 is detected in the proliferative zone, and NPR3 is predominantly expressed in hypertrophic chondrocytes^112^. Altered NPR3 expression may therefore have zone-specific effects that are not consistent with the constitutive loss of function phenotype. Further, in addition to its role as a ligand clearance receptor, NPR3 has been reported to function as a signaling receptor by modulating levels of cAMP and downstream PKA activity in the cranial placode^82^. It is therefore possible that NPR3 has additional roles in regulating long bone elongation besides CNP clearance. Regardless of this remaining uncertainty, our data support the hypothesis that natriuretic peptide signaling is an important regulator of vertebral skeletal proportion by modulating growth plate height and hypertrophic chondrocyte size^60^.

Several genes that appear in multiple of these skeletal proportion datasets do not have a known or clear role in cartilage development. Four of these genes (*Lmcd1, Nlgn3, Ttll9,* and *Zim1*) that we identified in the disproportionately elongated jerboa TV6 and jerboa metatarsals are also among differentially expressed genes in the mouse and rat limb proportion study^28^. While mutations in *Lmcd1* and *Nlgn3* are reported to cause abnormal bone mineral density and postnatal growth defects^57,113^, *Ttll9* and *Zim1* have no reported bone defects and have not been investigated in the growth cartilage.

Here, we have identified the cellular mechanisms of differential growth in the vertebral skeleton and candidate molecular regulators of proportion. Some of these are potentially specific to controlling proportion the vertebral skeleton while others suggest common mechanisms locally regulate growth rates throughout the skeleton. This opens avenues to focus attention on mutations that diversify proportion in natural populations, to identify *cis*-regulatory elements that modularize the timing and level of expression in different growth cartilages to achieve different growth rates, and to test the functions of genes not previously assigned a role in skeletal elongation.

## ACKNOWLEDGMENTS

We are grateful to all members of the Cooper laboratory for assisting in these experiments, especially Daniel Ochoa-Reyes and Sara Kamenski, and for discussing data and interpretations. We thank Jeffery Olgin at UC San Francisco for providing the *Npr3* knockout mice and Eric Chang, Saeed Jerban, and Qingbo Tang at the VA Medical Center in San Diego for use of their microCT scanner. We also thank Talia Moore, Terry Capellini, Matt Hilton, Richard Behringer, and Hopi Hoekstra for their mentorship and support of C.J.W. and Andrew McCulloch and Sam Ward for insightful discussion. This publication includes data generated at the UC San Diego IGM Genomics Center utilizing an Illumina NovaSeq X Plus that was purchased with funding from a National Institutes of Health SIG grant (#S10 OD026929). We thank Kristen Jepsen at the UCSD IGM for her help and guidance in designing and executing the sequencing experiments. This work was supported by the National Institutes of Health Ruth L. Kirschstein National Research Service Award (NRSA) Individual Postdoctoral Fellowship award number F32AR079923 to C.J.W. and the National Institutes of Arthritis, Skin, and Musculoskeletal Diseases award number R01AR075415 to K.L.C. We are also grateful to the Wu Tsai Human Performance Alliance and the Joe and Clara Tsai Foundation for supporting these studies.

## METHODS

### Experimental model and subject details

Jerboas were housed and bred as previously described^114^. CD-1 and C57BL/6 mice were obtained from Charles River Laboratories (MA, USA). NPR3 mutant mice (MGI:2158355) were obtained from Dr. Jeffery Olgin’s laboratory at University of California San Francisco. Male and female *Npr3*^-/-^ mice were initially crossed to C57BL/6 mice to expand the line. Due to the effect of NPR signaling on female ovarian biology and male penile function, both mutant and heterozygous animals were used for breeding to obtain mutant animals for study. All animal care and use protocols were approved by the Institutional Animal Care and Use Committee (IACUC) of the University of California San Diego.

### Skeletal preparations and image capture

Adult mice and jerboas were humanely euthanized, then carcasses were skinned and placed in an enclosed colony of dermestid beetles. Once the skeleton was cleared of tissue but joint articulations were still intact, skeletons were brushed off then frozen to kill surviving beetles/larvae. Skeletons were further cleaned by hand and brightened in a gentle 5% hydrogen peroxide solution.

Neonates were humanely euthanized, then carcasses were skinned and eviscerated and fixed in 95% ethanol overnight. Samples were stained overnight in cartilage staining solution (75% ethanol, 20% acetic acid, and 0.05% Alcian blue 8GX), rinsed in 95% ethanol, cleared overnight in 0.8% KOH, then stained again overnight in bone staining solution (0.005% Alizarin red S (Sigma-Aldrich, A5533) in 1% KOH). Specimens were cleared in 20% glycerol in 1% KOH until the entire axial skeletal morphology was visual to the eye. For imaging and storage, samples were processed through a series of glycerol in 1%KOH up to 100% glycerol. Protocol modified from Lim et al 2024^115^.

### Carcass preparation and uCT analysis

#### Carcass preparation

We performed transcardiac perfusion fixations to prepare entire carcasses for micro-computed tomography imaging. Animals were terminally anesthetized with a ketamine/xyaline solution. Once sufficient anesthesia was reached, we opened the rib cage to expose the heart and inserted a 21-gauge butterfly needle attached to a 50 cc syringe with warmed sterile saline into the left ventricle and cut a small incision into the right atrium to allow flow through. Saline was slowly flushed until the liver lost its red color and the solution flowed clear. The syringe was then exchanged for a 50 cc syringe containing freshly prepared 4% PFA, which was slowly injected until muscle twitching followed by limb stiffness was observed. Animals were then transitioned into 70% EtOH for storage. We collected >6 animals for each time point. Both males and females were collected at each time point for each species.

#### µCT image collection and analysis

CD1 mice and jerboas were scanned at 9 µm^3^ voxel resolution on a Skyscan 1076 MicroCT machine (50 kVp, 200 µA, 0.5 mm aluminum filter, 180° scan, Δ=0.8°). Specimens were supported during scanning using plastic pellets. Images were reconstructed using NRecon (Bruker, Belgium) with a smoothing factor of 1, ring artifact reduction factor of 6, beam hardening correction factor of 40%, and with a dynamic range from 0 to 0.11 attenuation units. Each reconstructed dataset was immediately viewed in Dataviewer to assess the quality of each scan. Due to a defect in the scanner that arose late in the experimental progress of this manuscript, some individuals were scanned twice (A/B numbered scans) so that all vertebrae could be measured without interference. NPR3 mice were scanned together on a GE eXplore 120 Preclinical micro-CT scanner (Waukesha, WI, USA) at 50 µm isotropic voxel size (100×100×165mm3 field of view, 60kV voltage, 32mA current, 0.5° rotation step).

Whole µCT files were opened for analysis in Bruker DataViewer software. Due to the size of the files, scans were always resized by a factor of 3. Skeletons were first oriented to the first sacral vertebra which is easily identifiable due to its unique morphology. The vertebral centrum was aligned in all axes before measuring the diaphyseal bone length in each element in the sagittal plane in sequence to the tail tip. The growth plates including the epiphyses were not included in measurements, because epiphyses have not formed by birth, and joint interzones cannot be detected by µCT.

To make a measurement, the cursor was drawn from the outside of the cranial end of the diaphysis in a straight line to the caudal end. Pixel intensities were measured over this distance and saved in an individual .csv file for each vertebra. The length of the vertebral diaphysis was determined to begin at the midpoint between the first pixel intensity minimum/maximum and to end at the midpoint of the last maximum/minimum. Pixels were converted to microns based on the scan resolution and resizing factor to determine the length of each element. Vertebral lengths were normalized to the naso-anal distance per each animal. Normalized vertebral lengths and relative change in vertebral length per week were visualized using ggplot2^116^.

### Growth cartilage tissue histology

#### Calcein pulse to measure growth rate

Mice and jerboas received an intraperitoneal injection of 15 mg/kg calcein. Tails were collected exactly 48 hours after injection and fixed overnight in cold 4% paraformaldehyde. Tails were then transitioned through a sucrose gradient into Optimal Cutting Temperature (OCT) media, dissected into two segments for sectioning: sacral 4 (S4) to TV2 and TV2 to TV8. Samples were then flash frozen in OCT media in block molds. Blocks were serially tape sectioned using Leica tape sectioning solution and CryoJane materials at 50 micron thickness. For imaging, slides were thawed and gently rinsed in PBS to remove OCT and tape residue. Sections were incubated with DAPI in PBS for 2 min then mounted in Fluoromount-G Mounting Medium. Sections were imaged on an inverted Olympus Fluoview 3000 laser scanning confocal microscope at 20x magnification. To capture the entire calcein front, and not pieces of the trabeculae, 10 optical sections were captured across 20 microns oriented around the center of the section. Optical sections were visualized in a maximum intensity projection and the distance between the calcein front and chondro-osseus junction, was quantified by spline interpolation using the InteredgeDistance Macro (Santosh Patnaik) in FIJI^117^ with default settings. Distances were averaged across three sections per individual for every growth plate and visualized in Graphpad Prism.

#### EdU Click-iT kit to quantify proliferation index

Mice and jerboas received an intraperitoneal injection of EdU per the manufacturer’s protocol and were collected exactly 2 hours after injection. Tails were fixed and prepared for cryosectioning as described above. Blocks were tape sectioned using Leica tape sectioning solution and CryoJane materials at 10 micron thickness. For imaging, slides were thawed and gently rinsed in PBS to remove OCT and tape residue. EdU was then detected using the EdU Click-iT protocol, and slides were mounted in Fluoromount-G Mounting Medium for imaging on an inverted Olympus Fluoview 3000 laser scanning confocal microscope. Three sections were imaged per individual for each growth plate.

Images were cropped to an ROI that captured the whole proliferative zone as determined by Hoechst co-stain and refined to the top/bottom-most EdU positive cell. Using FIJI, images were thresholded in each channel to create a mask. Watershed function was used to separate adjacent nuclei and ‘Analyze Particles’ function was then used to count each size-restricted shape. To calculate the proliferative index, the ratio of EdU positive cells to total number of cells per ROI was averaged across three sections for each individual. Results were visualized in Graphpad Prism.

#### Hematoxylin and Eosin staining for growth plate histology

Sections at 10 micron thickness were used for H&E staining following the University of Rochester Center for Musculoskeletal Research core protocol. Three sections per individual for each growth plate were imaged on an Olympus BX61 upright compound microscope. Spline interpolation was used to measure the height of the entire growth plate, hypertrophic, proliferative zone, and resting zone based on cell morphology. Average maximum cell heights were measured in the axis of elongation through the three largest cells with a clear nuclear profile and averaged across three sections. Results were visualized in Graphpad Prism.

### RNA sequencing and analysis

Tails were dissected in ice cold PBS and then incubated in Proteinase K for 10 minutes as described previously to remove tendons and connective tissue^29^. Tails were then washed in ice cold PBS and then TV1 and TV6 cranial growth plates were microdissected from the tail by cutting at the intervertebral disc and chondro-osseous junction. As much IVD tissue as possible was cut away but some annulus fibrosus remained in all samples. Three cranial growth plates per vertebral identity per species was pooled into RNAlater (Invitrogen) and equilibrated overnight at 4°C. Samples were removed from RNAlater and snap frozen in liquid nitrogen the next day and stored at −80°C or immediately processed for RNA extraction.

For tissue disruption, growth plates were ground into a powder using a mortar and pestle over liquid nitrogen then homogenized using the Qiagen QIAshredder system per the manufacturer’s protocol. RNA was extracted from the tissue homogenate using the Qiagen RNeasy Micro kit following the manufacturer’s protocol, including the on-column DNA digest. Initial RNA concentrations were recorded on a nanodrop before submission to the UCSD IGM core for tape station measurements. Only samples with RIN scores >8 were used for library preparation. UCSD IGM used Illumina mRNA Stranded Library kits to prepare libraries. Samples were run on PE100 lanes to get 25 million reads per sample on an Illumina NovaSeq S4 platform.

Quality control analysis and trimming of raw fastq files were performed using Trim Galore, a wrapper around Cutadapt^118^ and FastQC^119^. Trimmed reads were mapped to TOGA-annotated one-to-one and one-to-zero transcripts using STAR^120^, and read counts per gene were acquired by “--quantMode GeneCounts” option. STAR aligned gene counts were used to perform differential expression analysis between mouse and jerboa TV1 and TV6 with DESeq2^42^. To account for variable orthologous/transcript lengths between species we normalized gene lengths in the DESeq2 pipeline as described in Saxena et al., 2022^29^. Default DEseq2 settings were used to perform Principal Component Analysis (PCA) to identify variance associated with our tissue and species comparisons (n = 4 for all species/tissues). DESeq2 differential expression analysis was performed using the Wald test, and the DESeq function performed log_2_ fold-change shrinkage by default. We considered all differentially expressed genes with an adjusted p-value < 0.05 to be statistically significant in our analyses. R was used to perform all subsequent analyses and all generated code will be available on Zenodo.

The GO term enrichment analysis was performed using the clusterProfiler package^44,45^. We used the following settings: enrichGO function, org.Mm.eg.db mouse genome, SYMBOL keytype, BP (biological process) subontology (ont), all genes expressed across all growth plates in this analysis constituted the universe, and pValue cutoff was set at 0.05.

### Quantification and statistical analysis

To test the effect of sex on tail elongation, we performed a two-way ANOVA on mouse male and female individual relative tail lengths at each time point and found that there is no significant difference between male and female relative tail length. To test the effect of species on tail elongation, we performed Welch’s t-tests at each time point between mouse and jerboa and found that relative tail lengths are not significantly different from P0 to P14, and jerboa tails become significantly longer from P21 onwards.

We used the Shapiro-Wilk and Kolmogoroc-Smirnov tests to determine the normality of our histological measurement data and found that not all parameters are normally distributed, thus we performed an unequal variances t-test or Welch’s t-test to compare means between groups. We performed Welch’s t-test to compare cranial/caudal growth plates within species (i.e. mouse cranial TV1 vs. mouse cranial TV6) or growth plates of a single vertebrae (i.e. mouse cranial TV1 vs. mouse caudal TV1).

We used the GeneOverlap package^121^ in R to perform Fisher’s exact tests with Benjamini-Hochberg correction to compare the overlap between our 1,454 candidate genes and other datasets investigating skeletal proportion.

## SUPPLEMENTARY AFIGURES

**(Supplementary tables can be found separately in a series of Excel files)**

**Supplemental Figure 1.**
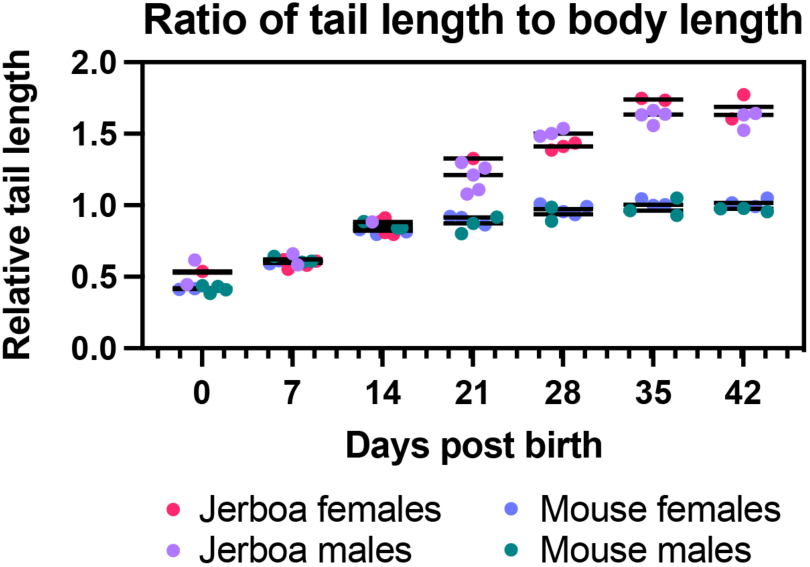
Tail lengths do not vary between male and female mice through development. A two-way ANOVA was used to compare the effect of sex on relative tail length in mice and jerboas and no significant difference was found.

**Supplemental Figure 2.**
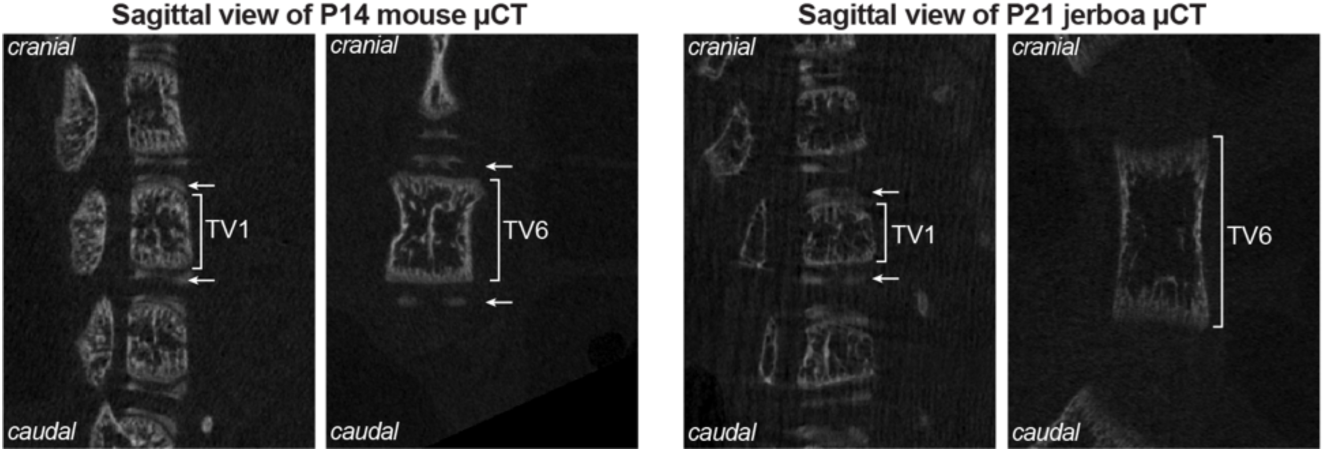
Endplate formation is evident in µCT scans of both species, and endplate ossification is delayed in jerboa TV6. Vertebral epiphyses (endplates, marked with arrows) form at the cranial and caudal ends of the vertebral diaphysis (brackets) by day P14 in mouse and in jerboa TV1 but not yet TV6 at P21.

**Supplemental Figure 3.**
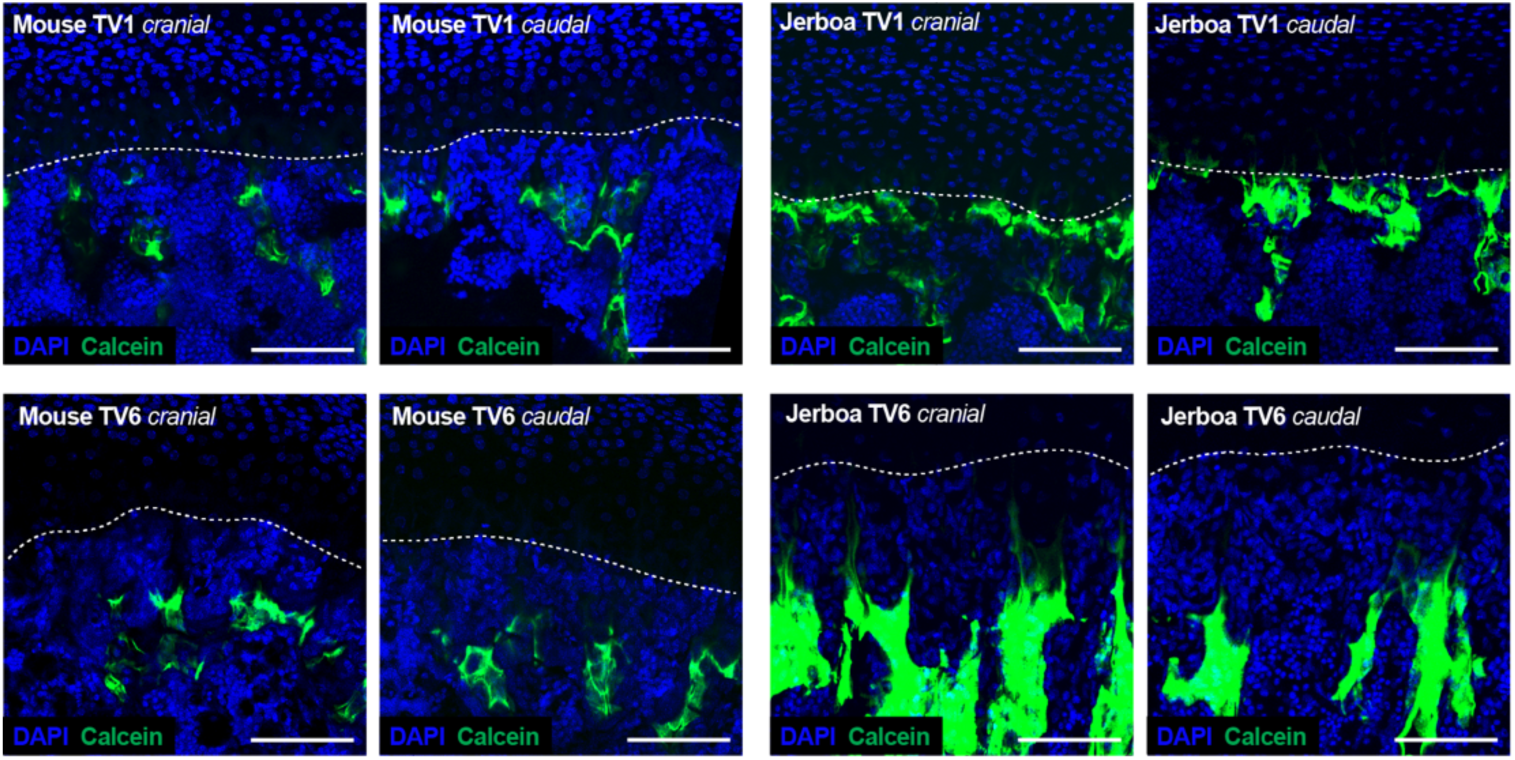
Visualization of daily growth using calcein dynamic histomorphology. Max intensity projections of calcein (green) labeling against nuclei (DAPI; blue). The chondro-osseus junction is identified with a white dotted line. Identity of each growth cartilage is indicated in the top left of every frame. Scale bar is 100µm.

**Supplemental Figure 4.**
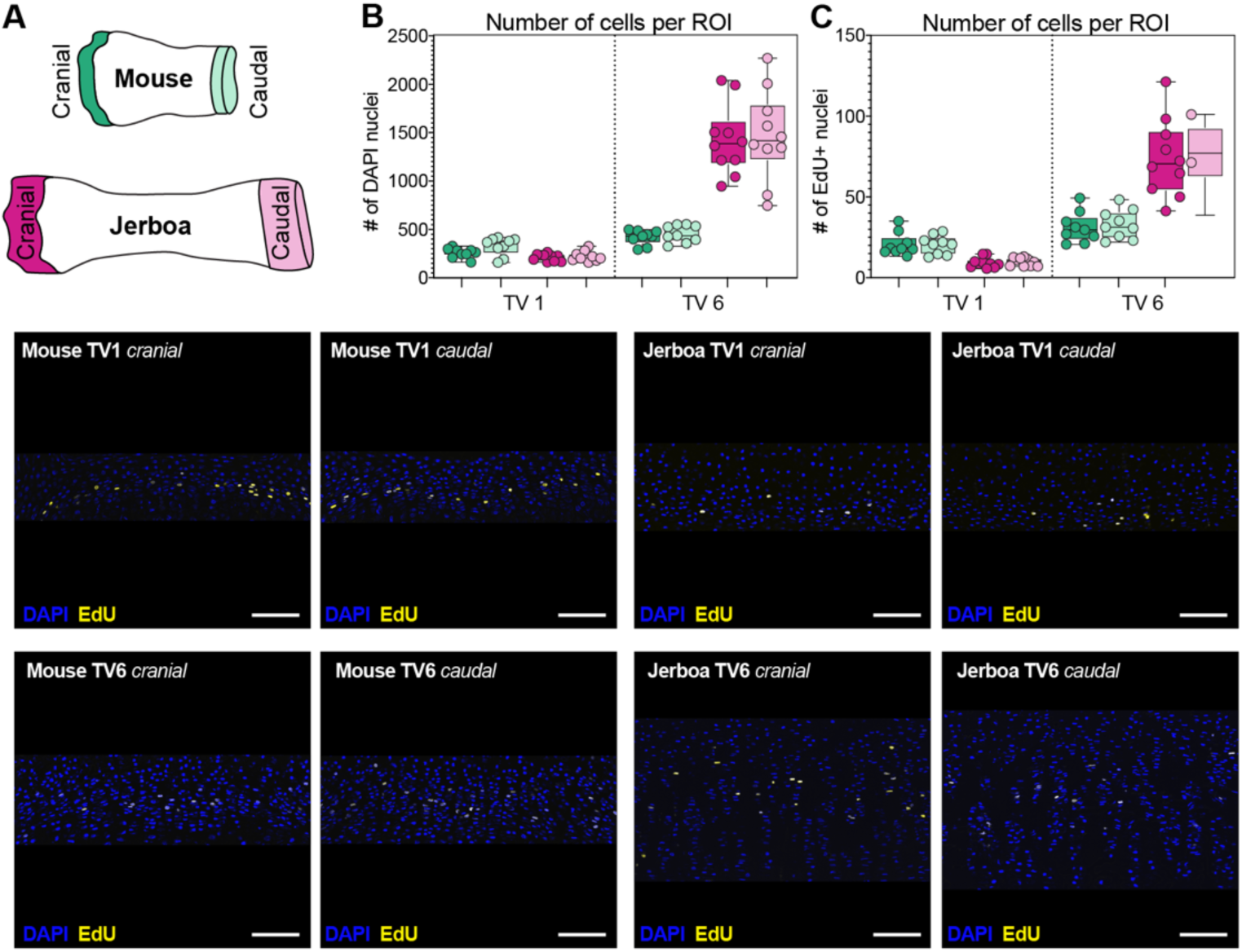
Visualization of EdU positive cells to determine proliferation index. (A) Drawing of mouse and jerboa vertebrae with corresponding color scheme. (B) Number of DAPI+ cells per ROI. (C) Number of EdU+ cells per ROI. (B-C) Each individual measured is indicated by a dot overlaying a box and whiskers plot. (D) ROI including whole proliferation zone. EdU (yellow) labeling against nuclei (DAPI; blue). Identity of each growth cartilage is indicated in the top left of every frame. Scale bar is 100 µm.

**Supplemental Figure 5.**
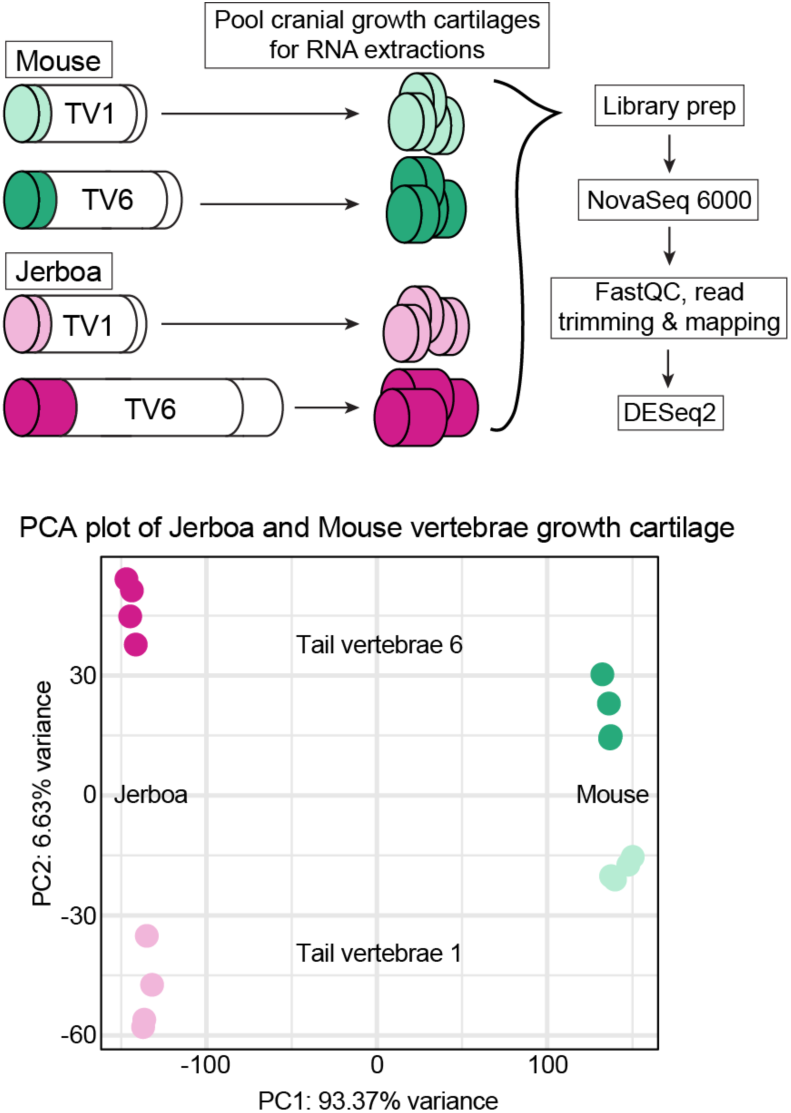
Experimental design of intersectional interspecies transcriptomics. (A) Schematic outlining experimental design for the RNA-sequencing experiments. (B) Principal components analysis of mouse and jerboa TV1 and TV6 replicates (n = 4).

## Notes

### Competing Interest Statement

The authors have declared no competing interest.

